# Spatial clustering reveals the impact of higher-order interactions in a diverse annual plant community

**DOI:** 10.1101/2025.07.14.664706

**Authors:** Theo L. Gibbs, Zachary J. Gold, Haylee Oyler, Jonathan M. Levine, Nathan J.B. Kraft

## Abstract

Spatial patterns are widespread in ecology, but their effects on species interactions remain unresolved, especially in diverse communities. In principle, the degree of spatial clustering could alter the concentration of higher-order interactions, which occur when one (or more) species modifies competition between two others. When species are well mixed, heterospecific neighbors have ample opportunity to modify a competitor’s interactions with other species. In contrast, species clustering can reduce the concentration of interspecific higher-order interactions. In a field experiment with annual grassland plants in California, we manipulated the spatial arrangement — but not the number or identity — of two competitors and measured how they jointly affected a focal individual. We found that focal plants produced more seeds when their competitors were clustered than when they were mixed. These results suggest that interspecific higher-order interactions generally had a stronger competitive (or weaker facilitative) effect than intraspecific ones. However, the effect of clustering varied across species. Larger differences in focal fecundity were correlated with competitors that had greater differences in size and/or functional traits between the spatial arrangements. Additionally, a competitive hierarchy among our study species predicted the effects of clustered versus mixed competitors on focal seed production. Altogether, our work suggests that the spatial arrangement of competitors changes the realized strength of competition in diverse plant communities by modifying the concentration of higher-order interactions. Given the extensive variation in spatial aggregation in plant communities, this mechanism is likely to be a powerful but underappreciated force shaping competition in nature.

**Significance Statement:** Plant species coexist in remarkably diverse assemblages throughout the world. Spatial patterns, including aggregation and intermixing, are also widespread in these communities. One potentially underappreciated mechanism that may structure the spatial dynamics of plant communities is interactions that uniquely occur in diverse systems, often called higher-order interactions. Here, we experimentally demonstrated that spatially mediated higher-order interactions operate among annual plants. These higher-order interactions, and their associated changes in competitor size and functional traits, were correlated with the competitive imbalance between competitors. Because both spatial aggregation and competitive hierarchies are widespread in nature, higher-order interactions emerging from their combination may be a more common driver of biodiversity patterns in plant communities than previously thought.

## Introduction

Spatial patterns and the aggregation of species are ubiquitous phenomena across ecosystems worldwide [1, 2, 3, 4, 5]. Accordingly, decades of effort have been devoted to understanding how patchy distributions arise, with ecological processes such as limited seed dispersal [6, 7, 1], heterogeneity of suitable habitat [8], and proximity to clustered mycorrhyzal mutualists [9, 10, 11, 12] all commonly invoked. However, how the resulting spatial patterns themselves shape the interactions between species in diverse communities is poorly understood.

Theoretical work has shown that aggregation modifies competitor dynamics [13, 14, 15, 16, 17, 18, 19, 5] largely because sessile organisms such as plants compete most strongly with others in their immediate neighborhood. If neighboring individuals are conspecifics, clustering can accentuate intraspecific competition and favor coexistence, a prediction supported by manipulative experiments [20, 21, 22, 23, 24]. Despite these advances, this body of work has focused largely on interactions between pairs of species. This is an important limitation because plant interactions, and the influence of spatial clustering, can be very different in diverse communities due to non-addivities or interaction modifiations [25, 26]. Diverse communities can therefore present a range of spatial patterns with no pairwise analogue, with as yet poorly explored implications for competitor dynamics.

One underappreciated way that spatial clustering might influence competition in diverse communities is through the operation of higher-order interactions [27, 28, 29, 30, 31, 32, 25, 33, 34]. A higher-order interaction occurs when one (or more) species modifies the pairwise interaction between two other species [30, 35, 36]. Critically, the concentration of particular kinds of higher-order interactions might depend on the patterns of spatial aggregation. When species are well mixed, heterospecific neighbors might modify a competitor’s interaction with another species, thereby resulting in an interspecific higher-order interaction (Fig. 1A). By contrast, clustered spatial patterns mean that competitors are surrounded by individuals of their own species. These conspecific neighbors might modify the interaction an individual has with its immediate competitors, resulting in an intraspecific higher-order interaction (Fig. 1A). Despite being the subject of much theoretical attention [32, 37, 38, 39, 40, 41], the relative strength of intraversus interspecific higher-order interactions in natural plant communities is largely unknown, even separate from the role of spatial patterns (but see [33, 26]).

**Figure 1.**
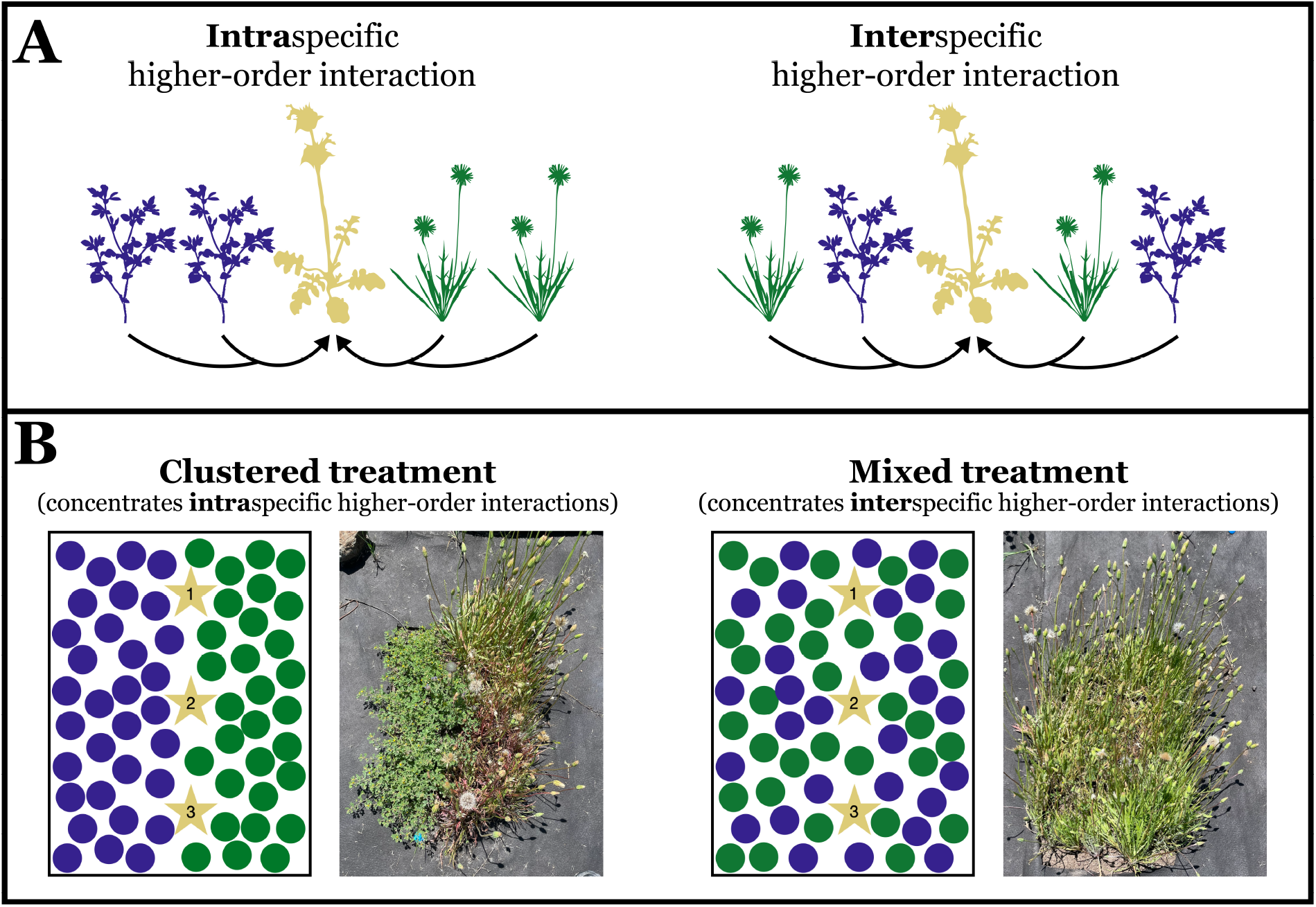
Types of interactions and experimental design. A) Intraspecific higher-order interactions occur when conspecific competitors (purple and green plants) modify each other’s interactions (arrows) with a focal individual (yellow plant). In contrast, when heterospecific neighbors change the competitive effect of one species on another, this generates an interspecific higherorder interaction. B) In the clustered competitor treatment (left cartoon and photograph), we spatially amplified the opportunity for intraspecific higher-order interactions to occur by planting the competitors (purple and green circles) on either side of the focal plants (yellow stars representing *Acmispon wrangelianus, Salvia columbariae*, and *Uropappus lindleyi*). In the mixed competitor treatment (right cartoon and photograph), the randomly interspersed competitors can interact and therefore result in more interspecific higher-order interactions. We ensured that the identity and densities of the competitors within a 10cm and 4cm radius around the focal individuals were the same in the clustered and mixed treatments, thereby holding pairwise competitive effects constant.

Perhaps more fundamentally, we lack mechanistic evidence for how higher-order interactions emerge from underlying processes [25, 35, 36], and therefore how they might respond to spatial aggregation in plant communities. Previous work [42, 43, 25] has hypothesized that neighborinduced changes in functional traits could allow one competitor to modify another’s effect on a third species. For example, a neighboring plant might alter a competitor’s rooting depth or leaf characteristics, and these changes can modify interactions with other species. At the same time, higher-order interactions could emerge simply when a competitor’s size is impacted by its neighbors. If a plant’s height or biomass is altered, then its competitive (or facilitative) effects on other species are likely to change as well [44, 45, 46, 47]. While differences in both size and functional traits due to competition are well-documented, [48, 49], their ability to subsequently modify the interaction between two other individuals has not been demonstrated. Doing so could potentially enhance our understanding of competition in diverse, spatially-structured plant communities.

Relatedly, interactions between species of unequal competitive rank could underlie spatially mediated higher-order interactions. Previous research on spatial clustering has shown that weaker competitors grow larger when clustered, while stronger competitors grow more poorly [14, 20, 21, 22, 50, 23, 51, 52, 49, 24]. Aggregation can dramatically reduce the performance of competitive dominants while only modestly amplifying the performance of competitive inferiors [20, 24], thus resulting in weaker overall competition. The role of a competitive hierarchy in spatially explicit competition may carry over to a multi-species context as well. From a higherorder interaction perspective, aggregation that minimizes interspecific higher-order interactions could promote weaker competition.

We conducted a field experiment manipulating the spatial pattern of competitors in an annual plant community in California to address three main questions: 1) How does the spatial aggregation of two plant competitors and, by inference, the relative strength of intraverus interspecific higher-order interactions influence the per capita fecundity of a third species? 2) How do changes in background competitor size and traits induced by clustering predict the effects of spatial aggregation on focal fecundity? 3) Can a competitive hierarchy explain how clustering affects focal plant performance? Our work aims to determine whether the combination of spatial aggregation and higher-order interactions could contribute to competitive dynamics in a diverse ecological community.

## Results

We performed an experiment manipulating the spatial clustering — but not the average number or identity — of two competitor species and measured their joint impact on a third focal species (1B). The experiment quantified how each of ten pairs of five background species (*Acmispon wrangelianus, Festuca microstachys, Plantago erecta, Salvia columbariae*, and *Uropappus lindleyi*) jointly affected the fecundity of a focal individual (one of three species: *A. wrangelianus, S. columbariae*, and *U. lindleyi*). In the clustered competitor treatment, we planted the two competitor species in a spatially aggregated fashion on either side of the focal plants to minimize the opportunity for these two competitors to interact, thereby dampening interspecific higher-order interactions while amplifying intraspecific ones (Fig. 1B). In the mixed competitor treatment, the two background competitors were intermixed, increasing the opportunity for interspecific higher-order interactions to occur and decreasing the concentration intraspecific higher-order interactions (Fig. 1B). Since the two spatial arrangements had a fixed density of competitors, the difference in focal plant seed production between them was designed to reveal the relative strength and direction of intra- to interspecific higher-order interactions.

We found that, when aggregating across the three focal species, focal individuals in our study produced more seeds with clustered competitors than with mixed competitors (Fig. 2). This reveals that interspecific higher-order interactions, accentuated in mixed plots, were more harmful (or less facilitative) than the intraspecific higher-order interactions accentuated in clustered plots. In our Bayesian linear models — with seed production log-normally distributed and fixed effects controlling for different focal identities and variation among plots (see Materials and Methods for more details) — we also found that the strength of this effect varied depending on the identity of the two background species (Fig. 2). Focal plants competing with *A. wrangelianus* in combination with any of the other four competitors produced markedly more seeds in clustered environments. The effect of the clustered treatment on seed production was positive for these four background competitors in 99.8% of draws from the posterior distribution (Fig. 2). The joint competitive effect of *S. columbariae* and *U. lindleyi* was also dampened when clustered, resulting in focal species producing more seeds in 99.9% of draws from the posterior. By contrast, the competitive effect of *F. microstachys* and *P. erecta* did not change appreciably in response to their spatial arrangement (37.6% of posterior draws were positive). Supplementary statistical analyses showed that the spatial clustering of competitors tended to increase seed production by approximately 28% aggregating across background species combinations (Supplementary Information). **Overall, competitors had a weaker effect and resulted in greater focal seed production when they were clustered compared to when they were mixed**. To evaluate potential mechanisms for the shifts in focal fecundity, we also evaluated a suite of traits for the background competitors in the two spatial arrangements. These metrics included two measures of plant size: plant height and aboveground biomass, both measured at peak biomass. We also used several widely measured plant functional traits, including specific leaf area (SLA), leaf dry matter content (LDMC), leaf carbon to nitrogen (C:N) ratio, and leaf carbon isotopic composition (*δ*^13^*C*, a measure of integrated water use efficiency) (see Materials and Methods and [53]). Consistent with our finding that competitor spatial arrangement influenced focal plant fecundity, background competitors exhibited different sizes and functional traits when they were clustered with conspecifics compared to when they were mixed with heterospecifics. However, the differences that emerged depended on the species. For example, three of five background competitors were taller when mixed with heterospecifics compared to when they were clustered: *F. michrostachys* (median [89th percentile Credible Interval]: 48.4cm [45.5cm, 51.2cm] mixed versus 44.3cm [42.3cm, 46.3cm] clustered), *S. columbariae* (18.0cm [16.1cm, 19.9cm] mixed versus 13.3cm [12.0cm, 14.7cm] clustered), and *U. lindleyi* (29.1cm [27.4cm, 30.7cm] mixed versus 26.9cm [25.7cm, 28.1cm] clustered) (Fig. **??**). For each of these three species, the magnitude of their height difference between treatments also depended on the other background competitor (Fig. **??**). These species did not significantly vary in biomass or SLA across treatments, but for a fourth competitor—*A. wrangelianus*—mixtures resulted in lower biomass (1.35g [1.02g, 1.69g] mixed versus 1.97g [1.74g, 2.21g] clustered) and smaller SLA (139.6cm^2^/g [131.9cm^2^/g, 147.3cm^2^/g] mixed versus 157.8cm^2^/g [152.4cm^2^/g, 163.1cm^2^/g] clustered) (Fig. **??**). Other traits, such as LDMC, leaf C:N ratio, and leaf carbon isotopic composition, did not substantially change between treatments for any species in our experiment (Figs. **??, ??, ??**, and **??**). When we combined the size and functional traits using principal components analysis (PCA) for each species (Fig. **??**), we found that spatial clustering affected the first principal component (PC1). Similar to focal plant fecundity, differences in PC1 across treatments were largest for species combinations that included *A. wrangeliaunus* as well as the *S. columbariae* and *U. lindleyi* combination.

**Figure 2.**
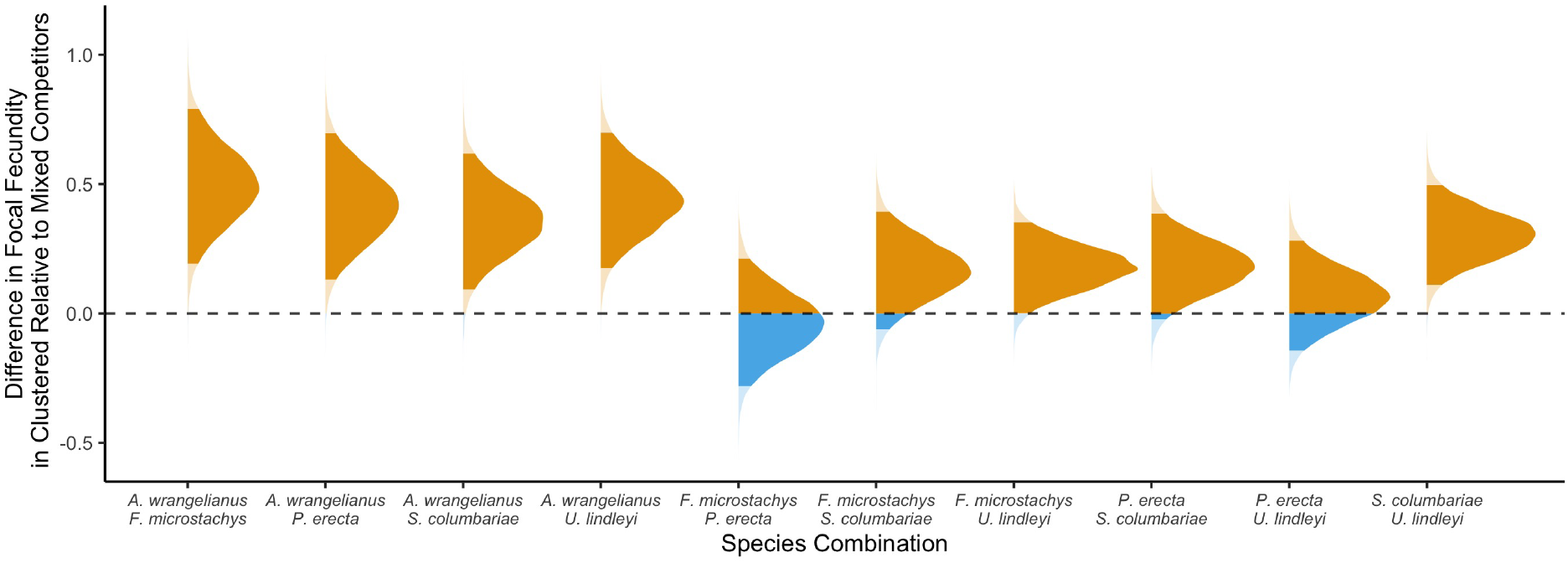
Differences in focal plant fecundity between clustered and mixed plots. Posterior densities for the estimated change in the fecundity of focal plants due to having clustered background competitors are shown across different background species combinations. Results are aggregated across focal species in a Bayesian linear model in which seed production was modeled with a log-normal distribution. The dashed vertical line denotes no change. The densities are colored orange when they are positive (indicating greater fitness in clustered plots), while blue colors indicate negative values. Colors are semi-transparent when outside the 95th percentile of the posterior density.

Indeed, background competitors with the greatest changes in response to clustering were those that resulted in the greatest differences in focal plant fecundity across spatial treatments (Fig. 3). More specifically, we found that across background competitor combinations, the degree of change in PC1 due to clustering was correlated with the degree to which focal plant fecundity responded to competitor spatial arrangement. We used a meta-analytic approach that accounted for uncertainty in both the independent and dependent variables to determine this correlation (see Materials and Methods for more statistical details), and the slope was positive in 90.2% of draws from the posterior distribution. **This result implies that the degree to which spatial structure influences the impact of inter- and intraspecific higher-order interactions can be predicted by the sensitivity of competitor size and functional traits to those spatial patterns**.

**Figure 3.**
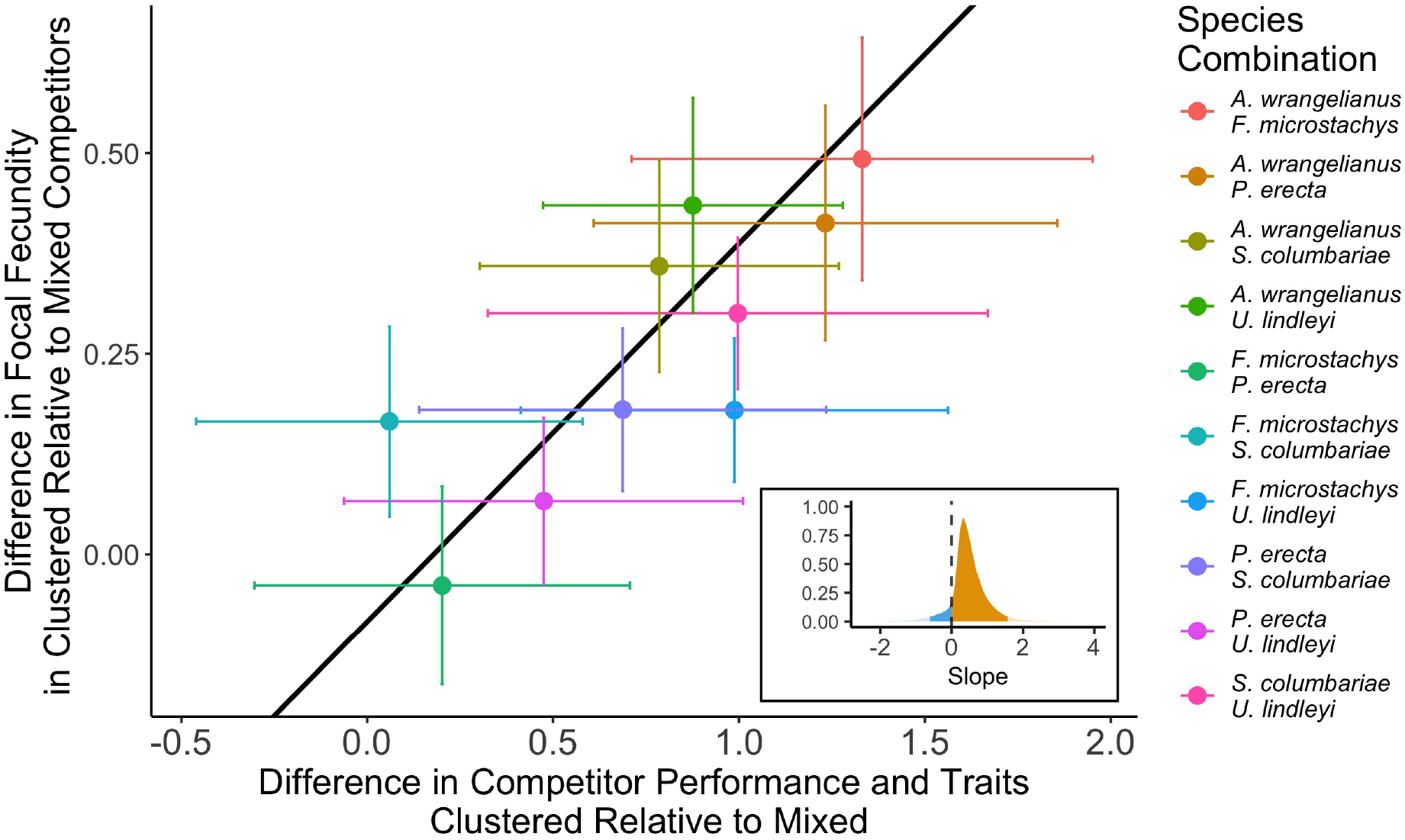
Species combinations with greater differences in size and functional traits resulted in greater differences in focal fecundity across treatments. Points are the average of the posterior densities for the estimated change in fecundity between clustered and mixed treatments plotted against the maximum estimated difference in the first principal component of the PCA for each species combination. Error bars denote one standard deviation in each quantity. The line is the best estimate of a Bayesian linear regression. Inset is the posterior distribution of the slopes from this regression. The dashed vertical line denotes a slope of zero. The density is colored orange when it is positive and blue when it is negative. Colors are semi-transparent when outside the 95th percentile.

Finally, we hypothesized that spatial aggregation might have particularly pronounced effects when the background competitor species had a large imbalance in their overall competitive impact. To evaluate this hypothesis, we quantified the competitive ability of each background competitor by averaging the reduction in fecundity caused by its presence across focal plant species and spatial treatments (see Materials and Methods for more details). Pairs of background competitors with greater imbalances in competitive ability produced larger differences in focal plant fecundity across the spatial treatments (Fig. 4A). The slope of the correlation between differences in fecundity and differences in competitive ability was positive in *≈* 99.9% of draws from the posterior distribution of a meta-analytic linear regression with uncertainty in the independent and dependent variables (see Materials and Methods for more details). For example, *A. wrangelianus* was a poor competitor compared to the other study species (Fig. **??**) and when it was one of the two background competitors, we found the largest differences in focal plant fecundity across the spatial arrangement treatment (Fig. 4). *S. columbariae* was the next weakest competitor while *U. lindleyi* had the largest competitive effects. Accordingly, the combination of *S. columbariae* and *U. lindleyi* as background competitors also generated large differences in focal fecundity across spatial treatment (Fig. 4). **Altogether, differences in the competitive ability among our study species were correlated with the differences in fecundity between clustered and mixed competitive environments, suggesting that competitive imbalances between species modulate the importance of spatial structure**.

**Figure 4.**
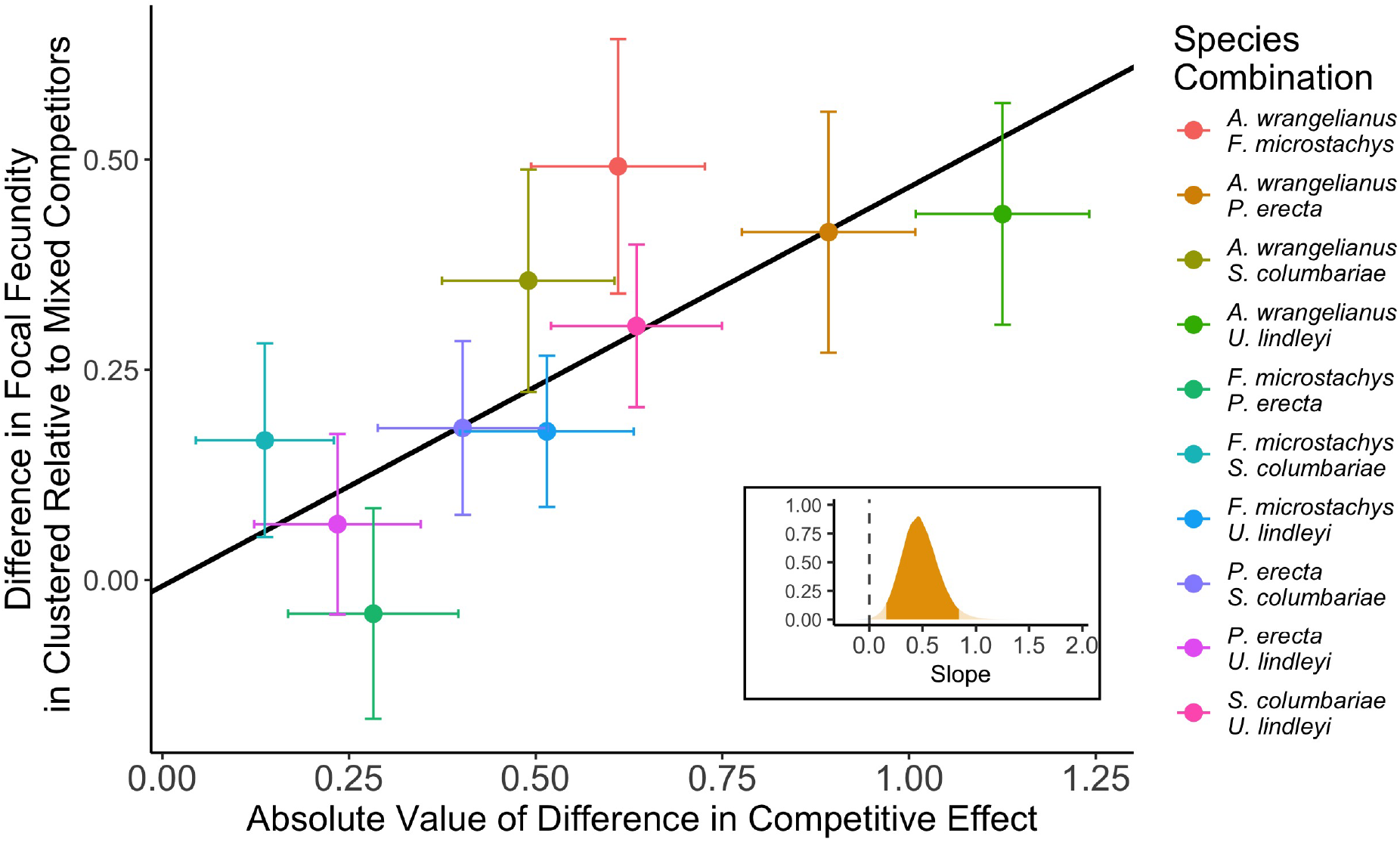
Competitive differences between species predicted the effects of spatial clustering. Points are the average of the posterior densities for the estimated difference in fecundity between clustered and mixed treatments plotted against the absolute value of the difference in competitive ability across species combinations (colors; same species codes as Fig. 2). Error bars denote one standard deviation in each quantity. The line is the best estimate of a Bayesian linear regression. Inset is the posterior distribution of the slopes from this regression. The dashed vertical line denotes a slope of zero. The density is colored orange when it is positive and blue when it is negative. Colors are semi-transparent when outside the 95th percentile.

## Discussion

An important challenge in community ecology is understanding how spatial patterns and species interactions reciprocally shape one another. Much previous research examining this problem focused on how species interactions produce spatial patterns [13, 14], but it is also possible for the aggregation of competitors to change their impact on other species [20, 54, 24]. Higher-order interactions (in which the pairwise interaction between two species is modified by the presence of another species) are particularly relevant in this context because they describe interactions that should be amplified or dampened by spatial clustering. Here, we found empirical support for the differential strength of intraspecific versus interspecfic higher-order interactions using a spatially explicit experimental design. Clustered competitors had weaker collective competitive effects on focal plants than mixed competitors, though results varied across species combinations (Question 1 from the Introduction). Given that clustering competitors should enhance the impact of intraspecific higher-order interactions and dampen interspecific ones, the weaker competitive effects observed in spatially clustered communities suggest that interspecific higher order interactions are more competitive (or less facilitative).

While previous work has hypothesized that changes in functional traits or plant growth might contribute to higher-order interactions [25], these connections have, to our knowledge, not been measured empirically. We found that the competitors with the greatest changes in size and functional traits in response to clustering also produced the largest changes in focal plant fecundity across spatial treatments. This result indicates that the magnitude of morphological change can be used to predict the strength of higher-order interactions (Question 2 from the Introduction). This finding is consistent with the notion that one background competitor impacts the growth and performance of the other, which in turn modifies its competitive effect on the focal. Indeed, some background species performed differently (reflected in the height and biomass differences shown in the Figs. **??** and **??**) depending on whether they were clustered or mixed with other species. Competition and spatial aggregation may have also introduced changes in the background competitors not directly related to growth and performance. For example, changes in functional traits such as SLA, leaf chemistry, or plant architecture (as measured by the canopy index) could reflect morphological or physiological differences that modify how these competitors interact with the focal plant species. While future work could study a wider range of traits, our results suggest that the effects of competition on the size and traits of background competitors propagated through to their interactions with other species. Thus, we provide experimental evidence that competition-induced changes in size and/or functional traits are linked to the strength of higher-order interactions, a first step towards a better mechanistic understanding of these interactions.

Differences in competitive ability between the two background competitors also predicted the effect of competitor clustering on focal plant fecundity and, by inference, the differing strengths of intraspecific and interspecific higher-order interactions (Question 3 from the Introduction). This result accords with past work which showed that, similar to our multispecies system, aggregating strong competitors in a pairwise context diminishes the impact of those competitors [20, 24]. In our study, competitive imbalances could have resulted in focal plants producing more seeds in the aggregated treatment for two non-mutually exclusive reasons. First, a focal plant with aggregated competitors could plastically respond to the spatial arrangement by, for example, preferentially uptaking resources from the side of the plot with the weaker species (as in [55, 56, 57]). Second, the stronger competitor could grow larger when mixed with a weaker one, thereby exerting more of a competitive impact on the focal plant. Our data on the background competitors supports both of these possibilities. In support of the first mechanism, *A. wrangelianus* was both the focal species most sensitive to spatial patterns (Fig. **??**) and showed the greatest plasticity in its growth form across treatments (Fig. **??**), which could potentially enable it to harvest light or other resources from the competitively weaker side of the plot. In support of the second mechanism, some of the strong competitors grew taller when mixed with the weakest competitor (Fig. **??**). Their larger size in the mixed plots could result in a greater competitive impact on focal performance compared to when strong intraspecific competition in the clustered treatment limited their growth. Future work could clarify the relative importance of different plastic responses and further disentangle the mechanisms linking changes in size or traits to species interactions.

Regardless of the underlying mechanism, the key role of competitive imbalances in explaining the effects of clustering on focal plant performance suggests that higher-order interactions may be common in nature. Competitive differences between species are pervasive [58, 53]; thus, diverse communities have ample opportunities for strong competitors to dampen their own effects when clustered and amplify their impact on other species when mixed with weaker competitors. These interactions are simpler than higher-order interactions that require unique phenotypic responses to competition to modify impacts on other species. Therefore, higher-order interactions might not be idiosyncratic phenomena arising from specific species combinations, but instead might be widespread in nature and reasonably predictable with a known competitive hierarchy.

Previously, determining the strength and effects of higher-order interactions in natural plant communities has proven challenging. Many approaches for measuring higher-order interactions, such as quantifying focal species performance as a function of two varying competitor densities [25, 33, 36], are labor intensive (due to the large number of required replicates) and challenging to assess statistically (due to uncertainty in model choice). Our approach circumvented these experimental challenges by spatially manipulating the opportunity for higher-order interactions to occur. Of course, the drawback is that this approach does not quantify the absolute magnitude of higher-order interactions. Rather, it reveals the relative sign and strength of interspecific to intraspecific higher-order interactions. Still, this information is useful for understanding how higher-order interactions might influence species coexistence. For example, the strong intraspecific higher-order interactions suggested by our results could make triplet coexistence more likely [39].

In general, our results illuminate a new way in which spatial structure can impact species interactions in diverse communities, underscoring the recent calls for more careful consideration of spatial patterns in community ecology [59, 60, 18, 5]. In natural plant communities, spatial aggregation is ubiquitous, emerging from varied mechanisms including limited seed dispersal, spatial resource heterogeneity, and even the clustering of mycorrhyzal mutualists. Competitive hierarchies are also a common ecological dynamic, suggesting that the different intra- versus interspecific higher-order interactions that arise from the combination of spatial aggregation and competitive imbalances may also operate in many plant communities. Our findings that differences in spatial aggregation patterns can shape competitor size, traits, and ultimately interactions in systems with more than two species thus provide an avenue for future progress. Ecologists should continue to explore the feedbacks between spatial patterns and higher-order interactions, connect higher-order interactions to a more mechanistic understanding of spatially-explicit competition, and explore how spatial patterns and higher-order interactions affect species coexistence.

## Materials and Methods

### Study Site

We established a field experiment in December 2023 at the University of California’s Sedgwick Reserve in northern Santa Barbara County, California, USA (34°40’ N, 120°00’ W). The site consists of heterogeneously distributed serpentine hummocks containing diverse assemblages of native annual plants. For this study, we selected five species representing a range of functional types and traits [53]: *Acmispon wrangelianus* (Fabaceae), *Festuca microstachys* (Poaceae), *Plantago erecta* (Plantaginaceae), *Salvia columbariae* (Lamiaceae) and *Uropappus lindleyi* (Asteraceae). This study was located in an approximately 500 m^2^ area that consists of finely textured serpentine soil and was cleared of vegetation, raked to facilitate planting, and fenced to exclude deer and gophers. Seeds for the experiment were collected from plants in prior experiments initially established from seed collections on the reserve.

### Experimental Design

We implemented the clustered versus mixed competitor treatments in 40 cm by 60 cm plots, with focal individuals of *A. wrangelianus, S. columbariae*, and *U. lindleyi* sown 15 cm apart (from each other and from the plot edge) along the center line of the plot’s long axis (Fig. 1B). Accounting for different viability among focal species, we planted enough seed to yield approximately 20 individuals at each focal location and subsequently thinned them to one individual after germination. We assigned these plots to receive background competitor seeds from one of the ten possible pairs of the five study species, seeded at 8 g*/*m^2^ in a clustered or mixed arrangement. *A. wrangelianus* was sown at a a reduced density of 6 g*/*m^2^ because of limited seed. We replicated each spatial arrangement treatment 15 times for each pair of background competitors, and established an additional treatment with no background competitors 15 times. We arranged the plots in a randomized block design throughout the experimental site, and the plots with background competitors reached densities roughly matching the natural density in the surrounding hummocks [58].

After germination, we manually removed individuals of all species that were not planted as part of our experiment. We also enforced the spatial pattern in the clustered plots by removing background competitors that emerged on the side of the plot in which they were not sown. To eliminate any resulting differences in competitor densities between the clustered and mixed treatments, we first censused the background competitor individuals within 4cm and 10cm from each focal plant and then thinned individuals in the mixed plots to remove any differences in plant density between treatments (see the Supplementary Information for full details). We also recorded the final number of background individuals, after thinning, for use in the statistical analyses. In the Supplementary Information, we provide a quantitative decomposition of how higher-order interactions contribute to differences in focal plant fecundity between clustered and mixed treatments.

### Data collection

In each plot, we randomly selected one individual of each competitor species to measure its end of season height, and we harvested the individual in order to measure aboveground biomass, specific leaf area (SLA), leaf dry matter content (LDMC), carbon to nitrogen (C:N) ratio, and leaf isotopic composition. Plants were wrapped in moist paper towels, sealed in plastic bags, and stored in a cooler for transport to the laboratory for measurements. For the leaf analyses, we selected several recently expanded, fully hardened leaves that were free of damage from each plant, weighed them, imaged them on a scanner at 600 dpi, and used ImageJ [61] to measure their area. We then dried the aboveground component of the plant, including all scanned leaves, to a constant mass at 60°C and weighed them to determine the leaf dry mass and total aboveground biomass. Finally, we ground several leaves from each plant to a fine powder for analysis conducted by the Center for Stable Isotope Biogeochemistry at the University of California, Berkeley. For *A. wrangelianus* and *P. erecta*, we also measured canopy index (as in [53]), which is the ratio of lateral to vertical growth. We defined lateral growth as the average of the longest axis (as viewed from above) and the axis perpendicular to the longest.

As plants senesced in May and June 2024, we measured focal plant fecundity in each plot. We did so by counting the number of seed pods for *A. wrangelianus*, flower clusters for *S. columbariae*, and seed heads for *U. lindleyi*, and then used statistical relationships between reproductive structure number and total plant seed production to predict per plant fecundity. We informed this relationship by measuring both variables on a sample of plants from the field (see Supplementary Information and Fig. **??** for full details).

### Statistical approach

We fit all statistical models following a Bayesian approach using the brms package [62] in R version 4.4.1 [63]. For each species combination, we fit a linear model to the seed production of focal individuals with fixed effects for focal species identity, the abundance of competing species as measured in our census, and the spatial arrangement treatment. Seeds were modeled as a log-normally distributed variable. In the Supplementary Information, we consider a suite of other statistical formulations and show that our main results are robust to modeling choices.

Subsequent statistical modeling tested how background competitors changed with the spatial arrangement treatment, and how this predicted the effect of that treatment on the focal plants’ seed production. We first used Bayesian linear models for each metric of size and functional traits to compare background competitors when they were clustered versus mixed (see Figs. **??**,**??, ??**, and **??**). Then, to determine the relationship between background competitor changes and focal plant fecundity across treatments, we ran a principal component analysis (PCA) using all of the measured traits in our study for each background competitor species. We took the first principal component (PC1) as an aggregate measure of variation in size and traits and determined how it was affected by the spatial arrangement treatment. Specifically, for each background competitor in each pairing, we ran a regression to determine the difference in PC1 in the clustered plots versus the mixed plots. Since a large change in just one competitor could result in a higher-order interaction, for each pair of background species we used the competitor with the larger difference in PC1 across the spatial treatments for further analyses (but see Fig. S17 for the same analyses with the average change of the two background species). We then used a meta-analytic approach to compare the competitor differences across spatial arrangements against focal fecundity differences across those treatments (see Supplementary Information for full details). This allowed us to account for error in both the dependent and independent variables.

To quantify the competitive ability of our study species, we used focal plant fecundity in plots with and without competitors to fit another Bayesian linear model, fit fixed effects for the presence of each competitor species on the fecundity of the focal plants, and did so separately for the clustered and mixed treatments. Summing together the competitive effect on focal plant fecundity across the clustered and mixed treatments yielded a single quantity describing how each background species impacted the focal plants. The difference in these quantities between two background species measured their competitive imbalance. We regressed the differences in focal species fecundity between spatial arrangement treatments against the absolute value of the differences between the competitive abilities of the background competitors following a metaanalytic approach. This analysis assesses the degree to which focal plant fecundity differences in the two treatments are larger when the background competitors are more imbalanced. See the Supplementary Information for a complete description of this statistical procedure.

## Supplementary Information

We provide additional analysis and figures.

## Acknowledgements

We acknowledge that the Land on which Sedgwick Reserve stands is part of the ancient homeland and traditional territory of the Chumash people. We thank Stephen Pacala and members of the Levin and Levine labs for helpful comments and discussion. We are grateful to the staff at Sedgwick Reserve for field coordination and to Carla D’Antonio for generously providing laboratory space at the University of California, Santa Barbara. We thank Jagveer Gill, Fidel Nigrete, Jessica Echarte, and Saba Saei for assistance preparing seeds for planting.

T.L.G. and Z.J.G. were each supported by the National Science Foundation Graduate Research Fellowship Program under Grant No. DGE-2444107. This work was supported by NSF Grant DEB-2022213, NSF Grant DEB-2022810, and the High Meadows Environmental Institute at Princeton University through the Mary and Randall Hack ‘69 Research Fund.

