## Supplementary Information for "Spatial clustering reveals the impact of higher-order interactions in a diverse annual plant community"

5 Figures 8

### 1 Experimental details

#### 1.1 Experimental setup

In December 2023, we cleared the 40cm by 60cm experimental plots of existing vegetation and tilled and raked by hand. To prevent sheeting during heavy rains, we installed straw wattles throughout the experimental area.

As described in the main text, each plot received background competitor seeds from two of the five species in a clustered or mixed arrangement. In clustered plots, we randomly switched whether each background competitor was assigned to the left or right side of the plot. We used a randomized block design to ensure that the results were not influenced by subtle differences in soil or microclimate. Each block received one plot of each treatment (clustered or mixed) for each background competitor species combination, and these plots were randomly assigned within the blocks. Blocks were then placed throughout the experimental site.

Following germination, we weeded plots for non-experimental plants throughout the season. In addition to removing plants that emerged from the seed bank, we further enforced the spatial patterns that we imposed at planting by weeding out heterospecific individuals that emerged on the wrong side of the clustered treatments (possibly due to either seeds getting shuffled prior to germination or the existing seed bank).

#### 1.2 Competitor census

To correct any differences in competitor density between the treatments introduced in the planting or weeding stages, we conducted two censuses of the neighborhood of the focal plants. On February 9th, 2024, we censused all germinants in a 4cm radius of every focal plant (irrespective of species identity). Within each background competitor combination and for each focal species, we matched clustered plots to mixed plots with a similar number of competitors in the census. On February 10th, we weeded out the difference in competitor individuals between the paired plots. From February 12-15, we censused all background individuals in a 10cm radius of every focal plant, recording the number and identity of the background competitors. Subsequently, we determined the difference between the average number of competitors for each background species in the two spatial treatments. We then weeded the plots to eliminate this difference within the 10cm radius.

#### 2 Quantitative description of spatial higher-order interactions

In the standard description of species interactions, the impact of one individual on another is described by the abundance of the competitor, its per capita competitive effect, and the distance separating the two competing individuals [1, 2]. Our experiment preserves the number and identity of competitors – as well as the distribution of distances between focal and competitor individuals – across the two spatial treatments. Therefore, phenomenological models that make these assumptions must invoke higher-order interactions to explain the differences in focal fecundity between the treatments.

More quantitatively, the fecundity of focal species $i$ (denoted $f_i$) is given by

$$f_i = r_i - \alpha_{ij} - \alpha_{ik} + \beta_{ijj} + \beta_{ikk} + \beta_{ijk} + \beta_{ikj} \tag{S1}$$

where $r_i$ is the number of seeds that species $i$ produces without competitors, $\alpha_{ij}$ measures the competitive impact of species $j$ on species $i$ and $\beta_{ijk}$ measures how the competitive effect of species $j$ on species $i$ changes when species $j$ competes with species $k$. Because each competitor in our experiment is planted at the same density across treatments and background combinations, we absorb any density-mediated effects into the competition parameters. Equation S1 is equivalent to a generalized Lotka-Volterra model for the dynamics of competing annual plants, but our argument in this section applies more generally to phenomenological models with the assumptions described in the preceding paragraph, including the Beverton-Holt model [3].

In plots without competitors, only the $r_i$ term is non-zero. In a clustered plot, the $\alpha_{ij}$ and $\alpha_{ik}$ terms both operate, describing the pairwise effect of each background species on the focal. Similarly, the $\beta_{ijj}$ and $\beta_{ikk}$ are present, because the background competitors compete with members of their own species. By contrast, the $\beta_{ijk}$ and $\beta_{ikj}$ terms are dampened because background competitors (species $i$ and species $j$) are spatially separated and hence do not interact. Only at the boundary of the plots in Fig. 1 of the main text can interspecific higher-order interactions occur in this treatment. In mixed plots, all of the terms in the decomposition of Equation S1 may be non-zero. The $\beta_{ijk}$ and $\beta_{ikj}$ terms are both present because background competitors now compete with members of the other species throughout the plot. On average, we expect the number of intraspecific higher-order interactions in a clustered plot to be evenly split between intra- and interspecific higher-order interactions in a mixed plot, neglecting any interspecific higher-order interactions at the boundary of split plots. Following this logic, the difference in fecundity between clustered and mixed plots directly reveals the relative strength of intra- to interspecific higher-order interactions, because the growth rates and pairwise interactions are the same in each treatment.

A more mechanistic understanding of the interactions between species may not need to employ higher-order interactions. For example, a spatially explicit but pairwise model of the biomass dynamics within the growing season could potentially produce higher-order interactions in a model of seed production between years and be a faithful representation of our experimental results. In nature, many higher-order interactions likely emerge from more basic pairwise processes rather than true many-body interactions.

### 3 Statistical approach

#### 3.1 Determining seed production

To determine the number of seeds produced by focal species, we counted the number of seed pods for each *A. wrangelianus* focal plant, flower clusters for *S. columbariae* focals, and seed heads for *U. lindleyi* focals. Independently, we collected data on the number of seeds per pod for *A. wrangelianus*, the number of seeds and the diameter of flower clusters for *S. columbariae*, and the number of seeds per head for *U. lindleyi*. Fig. S1 plots these relationships. For *A. wrangelianus* and *U. lindleyi*, we simply used the average number of seeds per pod or seed head to convert our field measurements of each focal to the number of seeds. For *S. columbariae*, we fit a linear model with seed number as the response variable and the squared diameter of the flower cluster as the independent variable, because we found this relationship fit the data better than the untransformed diameter. We used this linear relationship to determine the number of seeds using the diameter and number of flower clusters we measured in the field for each *S. columbariae* focal plant.

#### 3.2 Analysis of background competitor performance and functional traits

For each competitor species, we analyzed the effect of individuals being conspecifically clustered versus mixed with heterospecifics using a Bayesian linear model with spatial arrangement as a fixed effect. Results for performance metrics and functional traits are shown in Fig. S2 for clustered plots versus all mixed plots and in Fig. S3 for clustered plots versus mixtures with each heterospecific competitor (including raw data). We show the same data for the chemical and isotopic data in Fig. S4 and Fig. S5. In the main text, we only consider two of these chemical/isotopic traits chosen for their strong connection to biological function. These are the carbon to nitrogen (C:N) ratio, which is a widely used metric for the balance between two critical nutrients, and leaf carbon isotopic composition ($\delta^{13}C$), a measure of integrated water use efficiency. Since we were only able to measure canopy index in *A. wrangelianus* and *P. erecta*, the results from those models are shown separately in Fig. S6. The other species did not have substantial changes in their lateral growth. We also separately show model results for competitor heights when grown with *A. wrangelianus* as mentioned in the Discussion of the main text (Fig. S7).

To reduce dimensionality and compare changes across species, we subsequently compiled all of the performance metrics and functional traits into a principal components analysis (PCA) for each of the background competitor species (Fig. S8). We then used the first principal component to analyze how differences in competitor performance and traits correlated with differences in focal fecundity between the spatial arrangements, as described in the main text.

##### 3.3 Computing competitive effects

To quantify competitive effects described in the main text, we fit a Bayesian linear model to the fecundity data from plots with and without competitors. Specifically, we fit a fixed effect for focal species identity and then effects for the presence or absence of each competitor in each spatial arrangement. For example, we used the model to estimate the reduction in fecundity due to *A. wrangelianus* when it was clustered or mixed separately. To measure the total competitive effect of *A. wrangelianus*, we summed its effect in each spatial treatment. Importantly, this metric does not contain any information about the spatial arrangement of competitors or the identity of the other competitors. Fig. S9 plots the posterior distributions of the competitive effects across species. As described in the main text, *A. wrangelianus* has the weakest competitive effect, while *U. lindleyi* has the strongest. *S. columbariae* has the second weakest competitive effect, followed closely by *F. microstachys*. *P. erecta* was the second strongest competitor.

##### 3.4 Meta-analysis approach

To relate the effect of the spatial arrangements on focal fecundity to our analyses of performance/traits or competitive effects, we took a meta-analytic approach. This approach allowed us to use separate models for focal fecundity and the other analyses while propagating error. Specifically, we computed the mean and standard deviation of the posterior distribution of the focal seed production differences (as the dependent variable) and the mean and standard deviation of the posterior distribution of the independent variable (maximum change in phenotype as quantified by the first principal component or the absolute difference in competitive effect). We modeled the change in seed production using a Gaussian distribution and we incorporated uncertainty in the independent variable through measurement error. See the code at https://github.com/theogibbs/SplitMixed for further details.

### 4 Results from alternative statistical formulations

In the following subsections, we describe results from alternative statistical formulations and show that they are qualitatively consistent with those presented in the main text.

#### 4.1 Pooling across background combinations

To assess how the spatial arrangement of competitors influenced seed production across background species combinations, we fit a Bayesian linear model to the entirety of the fecundity data. Specifically, we fit fixed effects for the focal species identity, the background competitor species combination, and the spatial arrangement. In the model, we did not control for random variation in the background competitor abundances, because their per-capita effect could vary depending on the identity of the background competitors. As reported in the main text, focal species in clustered plots produced around $28\%$ more seeds than those in mixed plots (see Fig. S10 for the full posterior distribution which was positive for $> 99.9\%$ of draws).

#### 4.2 Effect of clustering on each focal species individually

Instead of including focal species identity as a fixed effect in our modeling framework, we also tested fitting different models for every focal species. Fig. S11 displays these results. Broadly, we see the same qualitative patterns within focal species as those we observed in Fig. 2 of the main text. As expected because these models are fit to less data, the inferred effect of the spatial arrangements are more variable. Nonetheless, focal species seed production in background competitor combinations that included *A. wrangelianus* still responded reasonably strongly even at this more granular level of detail (Fig. S11). Overall, we found that *A. wrangelianus* focal individuals had the strongest response to spatial clustering (positive in $\approx 88.6\%$ of posterior draws across background competitors), while *S. columbariae* and *U. lindleyi* produced more seeds in clustered environments in $\approx 74.1\%$ and $\approx 79.5\%$ of posterior draws respectively. Additionally, the inferred effect of spatial arrangement on *S. columbariae* fecundity was tightly concentrated when it competed with the several background combinations. This is because seed production was dramatically reduced in these competitive environments and *S. columbariae* focal individuals produced only a single flower cluster with minimal variation in diameter. Fitting a Poisson distribution to counts of the number of flower clusters from these competitor combinations produced more sensible posterior densities (Fig. S12), but these fits also discarded any information about the size of the flower cluster, which influences the number of seeds (Fig. S1). In either the continuous or discrete approach, however, we did not infer a strong effect of spatial arrangement in these background competitor combinations for *S. columbariae* focal individuals. Overall, we used a continuous approach in the main text because it is clearly appropriate for the other focal species and also for *S. columbariae* individuals in the other background competitor combinations.

We also modeled the identity of the focal species as a random effect and found qualitatively similar results for the difference in seed production (Fig. S13). Last, we included a random effect for the identity of the plot in the model used to generate Fig. 2 of the main text to control for spatial variation across the experimental area. Results from this regression were qualitatively the same as those presented in the main text (Fig. S14).

##### 4.3 Fitting a population dynamic model to fecundity data

Throughout our statistical approach thus far, we fit linear terms to account for the random variation in abundance (except when we did not fit any terms to account for this variation). The effect of competitor abundance on focal species fecundity, however, is often nonlinear [3]. In this section, we fit the Beverton-Holt model to our seed production data and show that our statistical results hold. Specifically, we fit the following relationship between seeds and competitor abundance:

$$f_i = \frac{r_i}{1 + \sum_j \alpha_{ij}^{clustered} N_j + \sum_j \alpha_{ij}^{mixed} N_j} \tag{S2}$$

where $f_i$ are the number of seeds and $r_i$ is the intrinsic seed production in the absence of competition as in Eq. S1. Further, $N_j$ is the abundance of competitors of species $j$ as measured by our census, $\alpha_{ij}^{clustered}$ is the competition coefficient measuring the impact of species $j$ on species $i$ in a clustered spatial arrangement, while $\alpha_{ij}^{mixed}$ is the same competition coefficient in a mixed environment. In this model, the impact of competition is nonlinear – as more individuals of any species are present in the neighborhood of the focal plant, their per capita joint competitive effect decreases. We follow a Bayesian nonlinear regression approach to fit this model. Specifically, we fit the growth rates $r_i$ and the competition coefficients $\alpha_{ij}^{clustered}$ and $\alpha_{ij}^{mixed}$ to all the data we collected in our experiment, including plots without competitors. Rather than compare the average change in seed production due to spatial arrangement, we now quantify the effect of the two spatial treatments by comparing the two different inferred competition coefficients. If the per-capita effect of competition for a given background competitor is the same, then the spatial arrangement did not change the strength of competition. As in our other statistical approaches, the effect of the spatial arrangement of competitors depended on the identity of the focal and background species (Fig. S15). However, aggregating across background competitor combinations, we still found strong support for the result that focal individuals suffered less from competition in clustered environments (Fig. S16). We quantified the effect of the spatial treatment on species $i$ across background competitors through the sum of log ratios: $effect_i = \sum_j \log\left(\frac{\alpha_{ij}^{mixed}}{\alpha_{ij}^{clustered}}\right)$. This quantity was positive (indicating stronger competition in mixed plots as in the main text) in $> 99.9\%$ of draws from the posterior distribution for *A. wrangelianus*, $\approx 85.3\%$ of draws for *S. columbariae* and $\approx 98.6\%$ of draws for *U. lindleyi*. Overall, we found that the elevated seed production we observed in clustered competitors is not due to this form of nonlinearity in the competitive dynamics between annual plants.

##### 4.4 Average difference in competitor performance and traits

In the main text, we analyze the effect of changes to the performance and traits of background competitors across spatial arrangements on differences in focal fecundity. Since we measured both background competitor species in every two species combination, we used the species with the maximum median difference in the first principal component between spatial arrangements for our analysis. For a higher-order interaction to occur, only one competitor needs to be modified such that its interactions change; thus, the maximum difference in performance and traits allows us to evaluate the competitor which might be most responsible for any higher-order interactions that occur. However, we also considered the average between the two competitors in their change in performance and traits. The same relationship holds, albeit slightly weaker ($87\%$ of posterior draws are positive), and with slight changes in some species combinations (Fig. S17).

#### 5 Figures

See the GitHub repository at https://github.com/theogibbs/SplitMixed for data and the code to run analyses and generate figures.

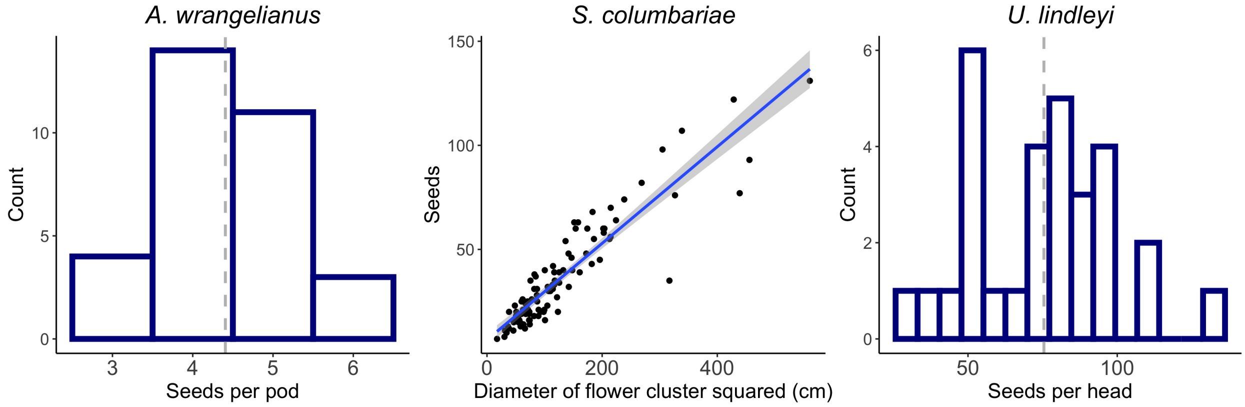

Figure S1. Histograms of the number of seeds per pods for *A. wrangeliaunus* and per seed head for *U. lindleyi*. The dashed lines show mean values. For *S. columbariae*, we plot the number of seeds against the squared diameter of flower clusters measured from a sample of flower clusters from background competitors. The blue line is a linear regression.

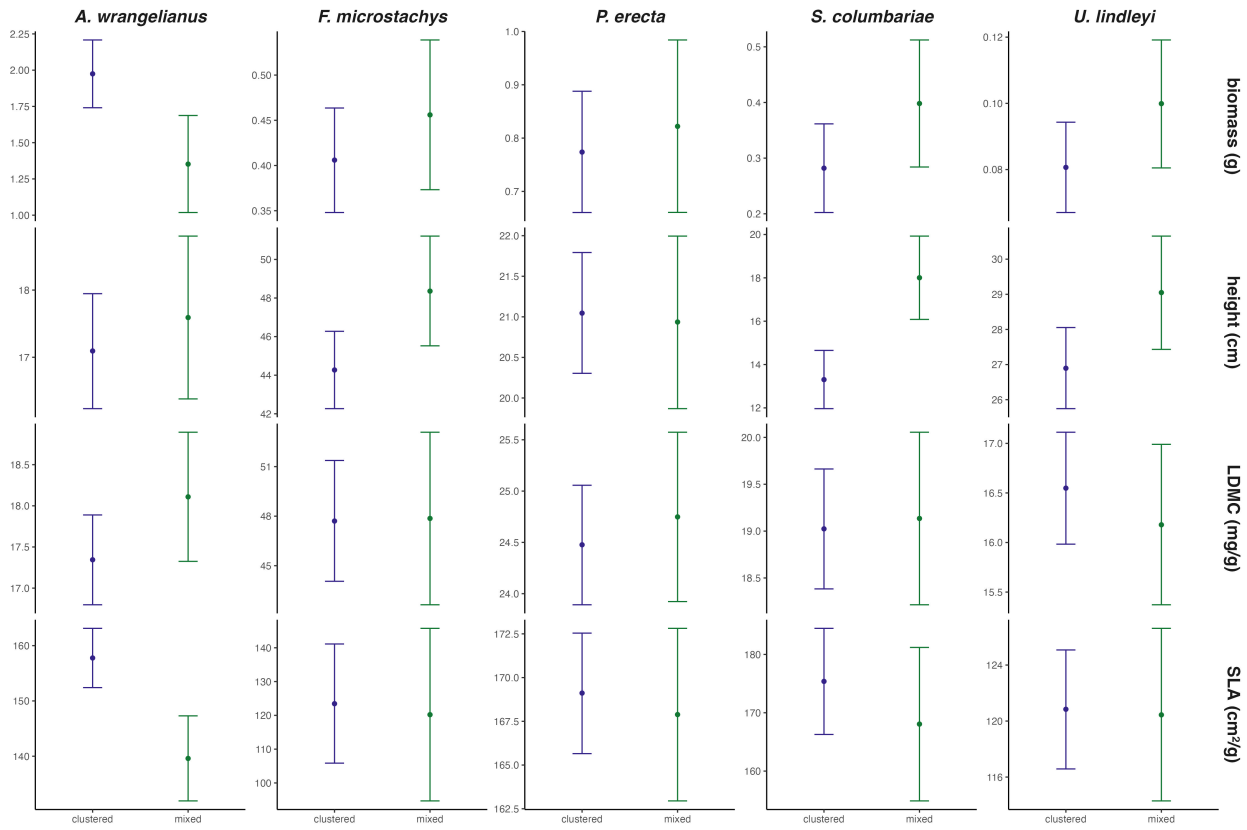

Figure S2. Background competitor performance metrics and functional traits in clustered versus mixed plots. Columns are species and rows are different metrics. Points are medians from the posterior and error bars are the $89\%$ credible intervals.

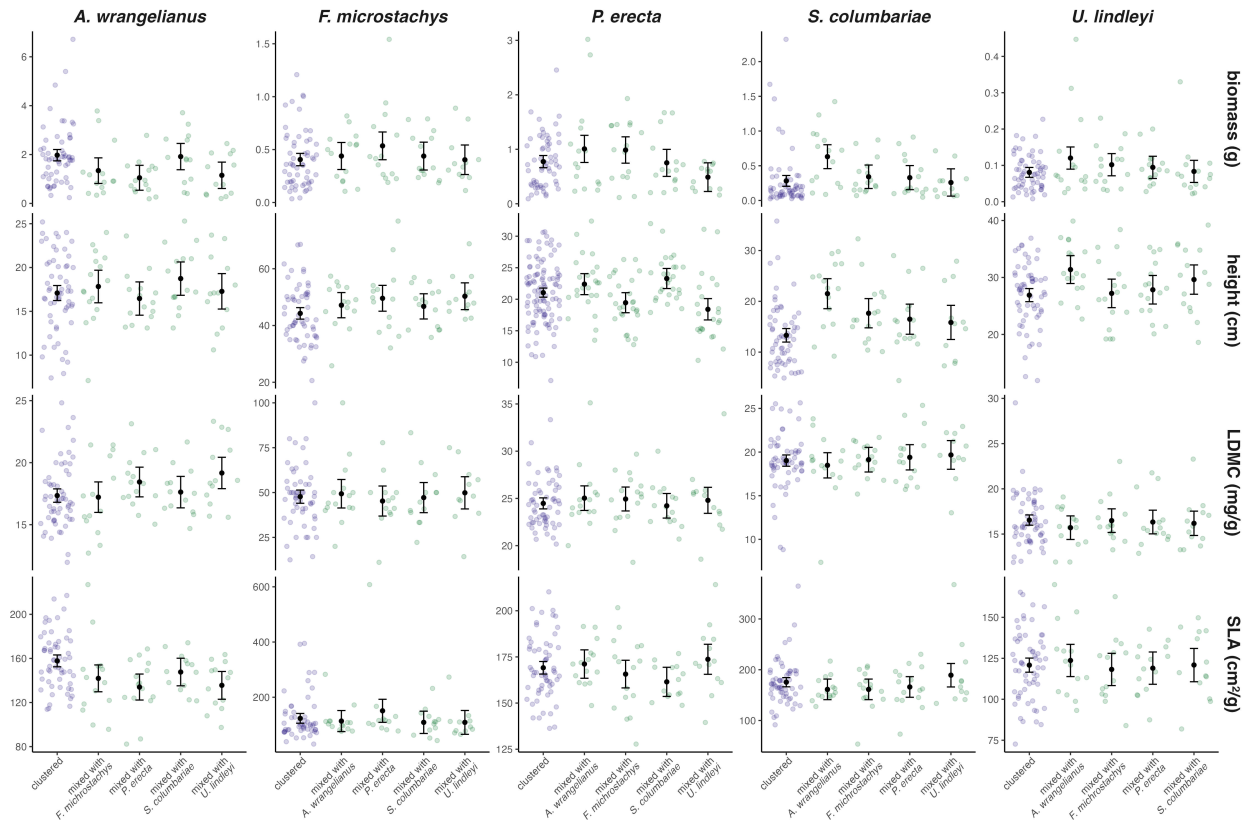

Figure S3. Performance and functional traits across all species combinations. Columns are background competitor species and rows are the different metrics. Points are medians from the posterior and error bars are the $89\%$ credible intervals. Circles are raw data.

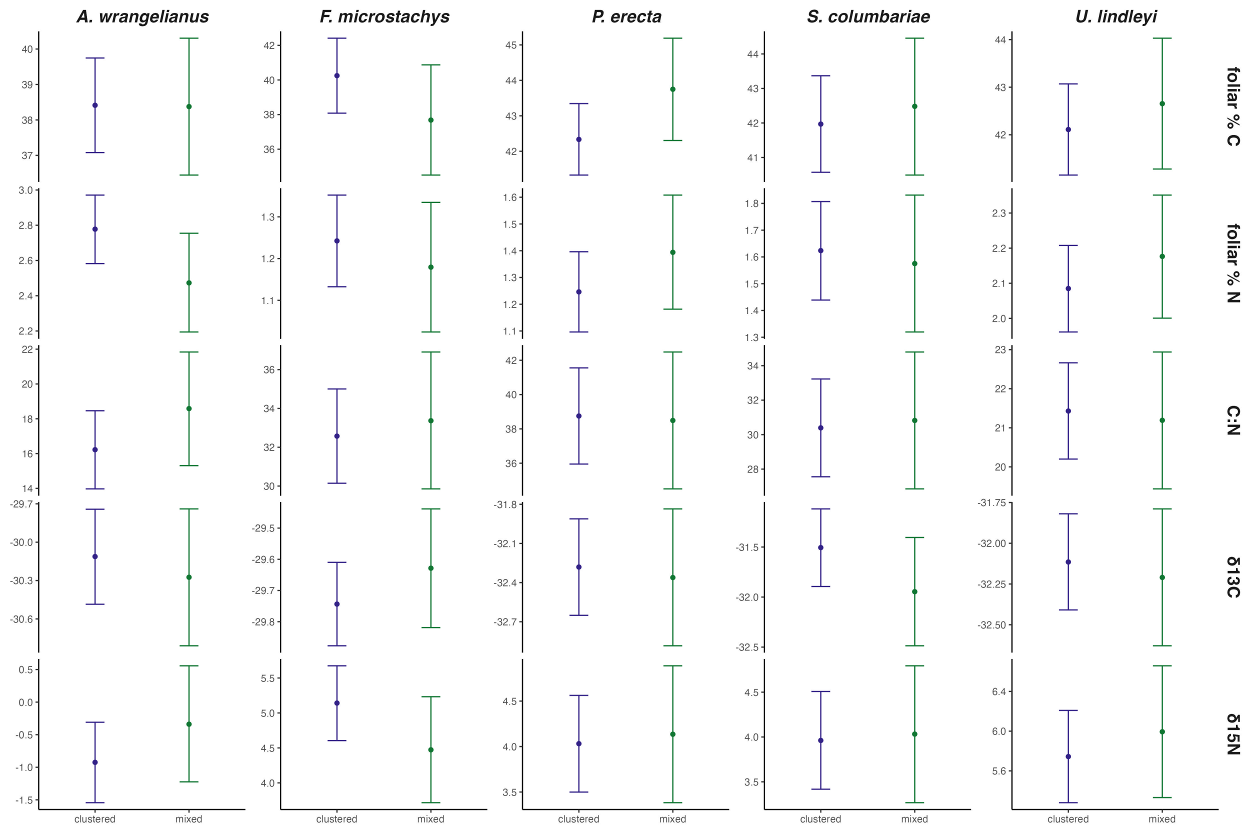

Figure S4. Background competitor chemical and isotopic traits in clustered versus mixed plots. Columns are species and rows are different metrics. Points are medians from the posterior and error bars are the $89\%$ credible intervals.

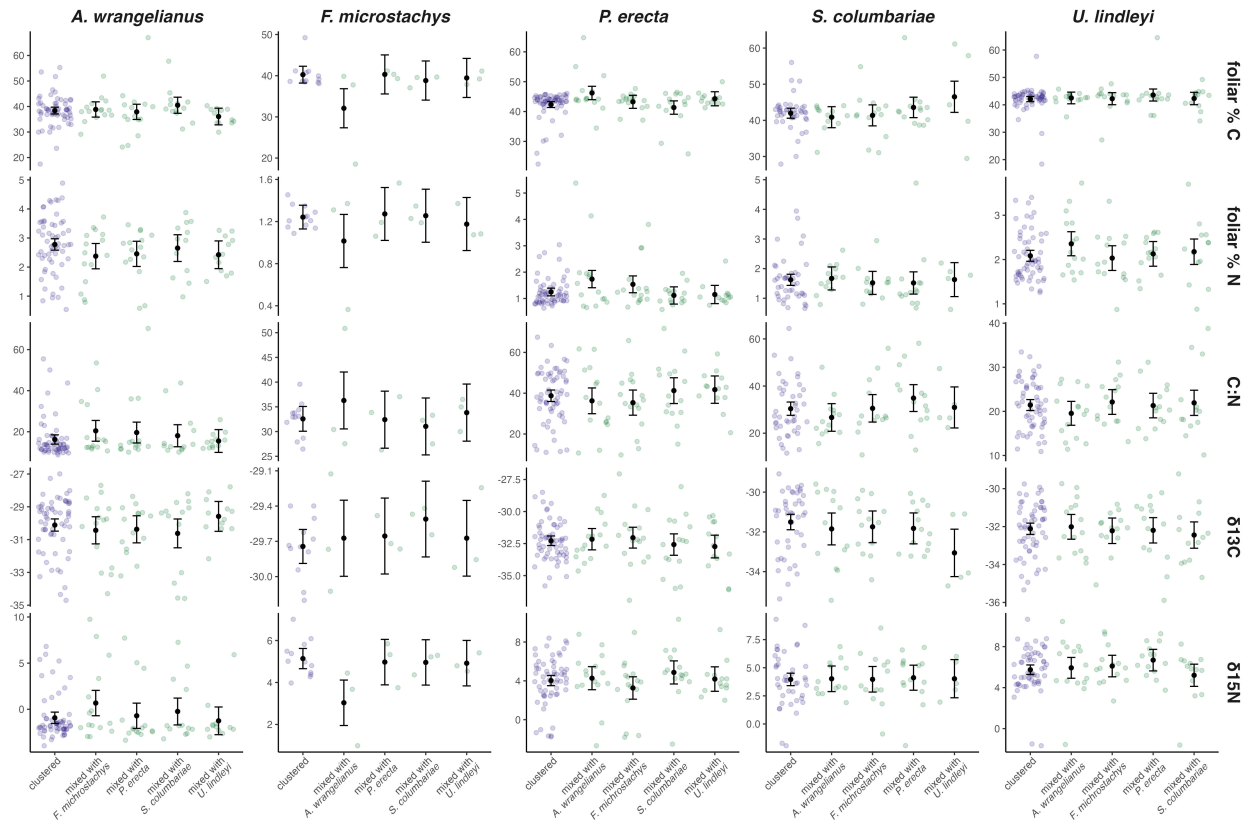

Figure S5. Chemical and isotopic traits across all species combinations. Columns are background competitor species and rows are the different metrics. Points are medians from the posterior and error bars are the $89\%$ credible intervals. Circles are raw data.

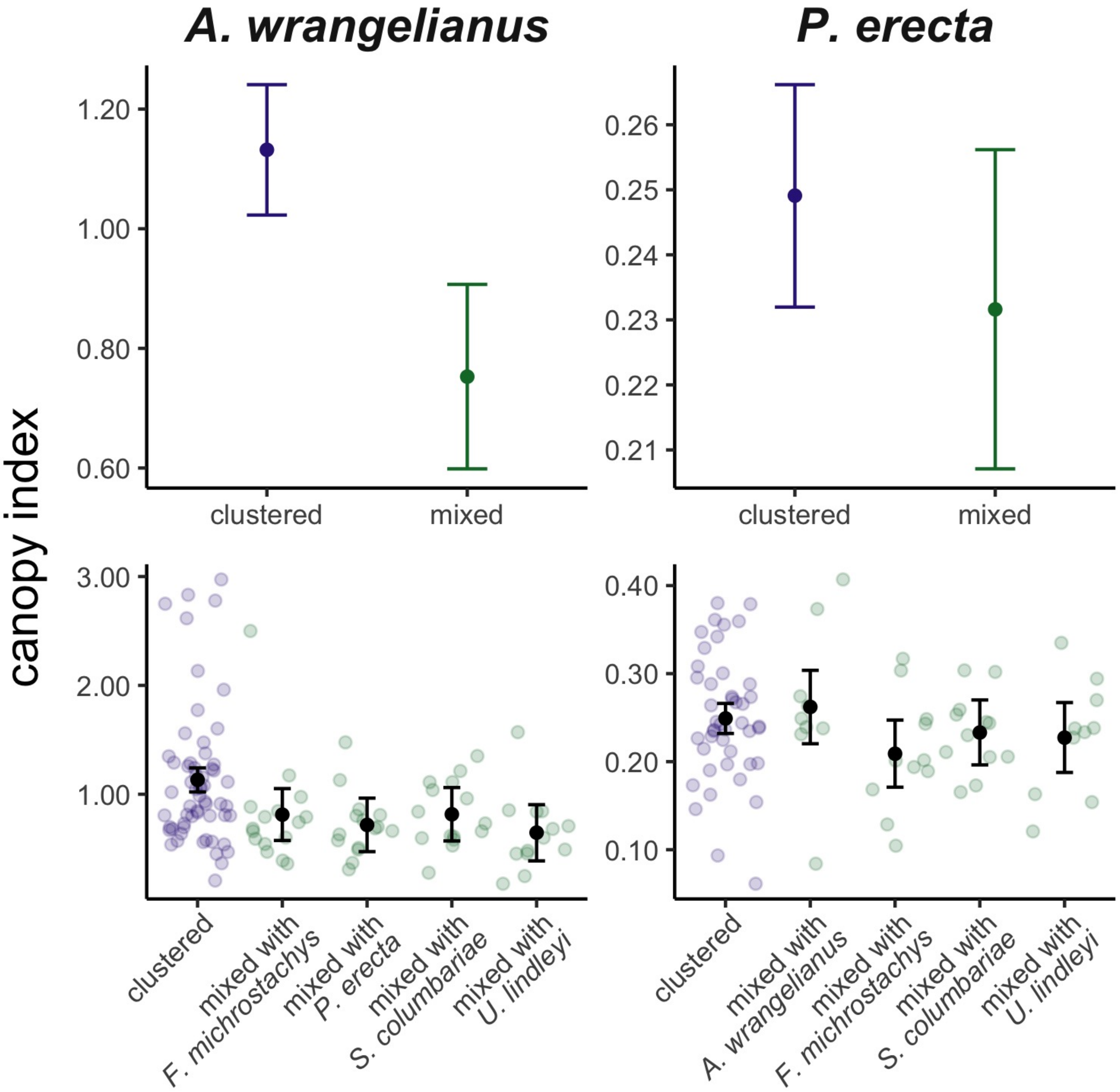

Figure S6. Canopy index for the two background competitors measured. The top row shows model results for clustered versus all mixed plots while the bottom row separates the mixed plots by species combination. All points are medians from the posterior and error bars are $89\%$ credible intervals. Circles are raw data. Canopy index was defined as the ratio of lateral to vertical growth, with lateral growth as the average of the length of the longest axis as viewed from above and the length of the axis perpendicular to the longest.

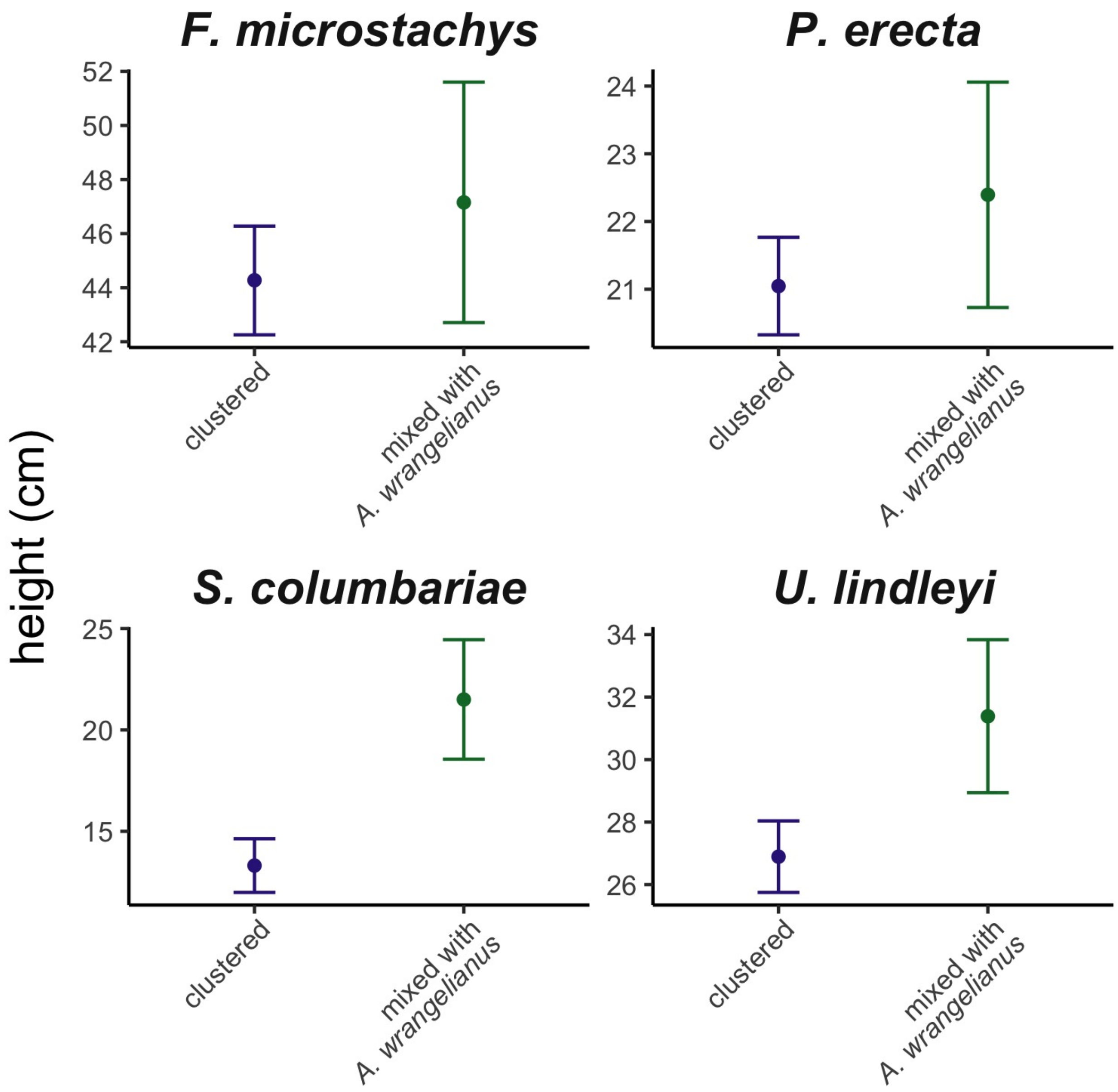

Figure S7. Competitor height with *A. wrangeliaunus* versus clustered. Model results show medians and $89\%$ credible intervals from the posterior. *A. wrangeliaunus* was the weakest competitor, and *S. columbariae* and *U. lindleyi* both show increased height when mixed with it as compared to when they were clustered with themselves or mixed with other heterospecific competitors (from Fig. S3).

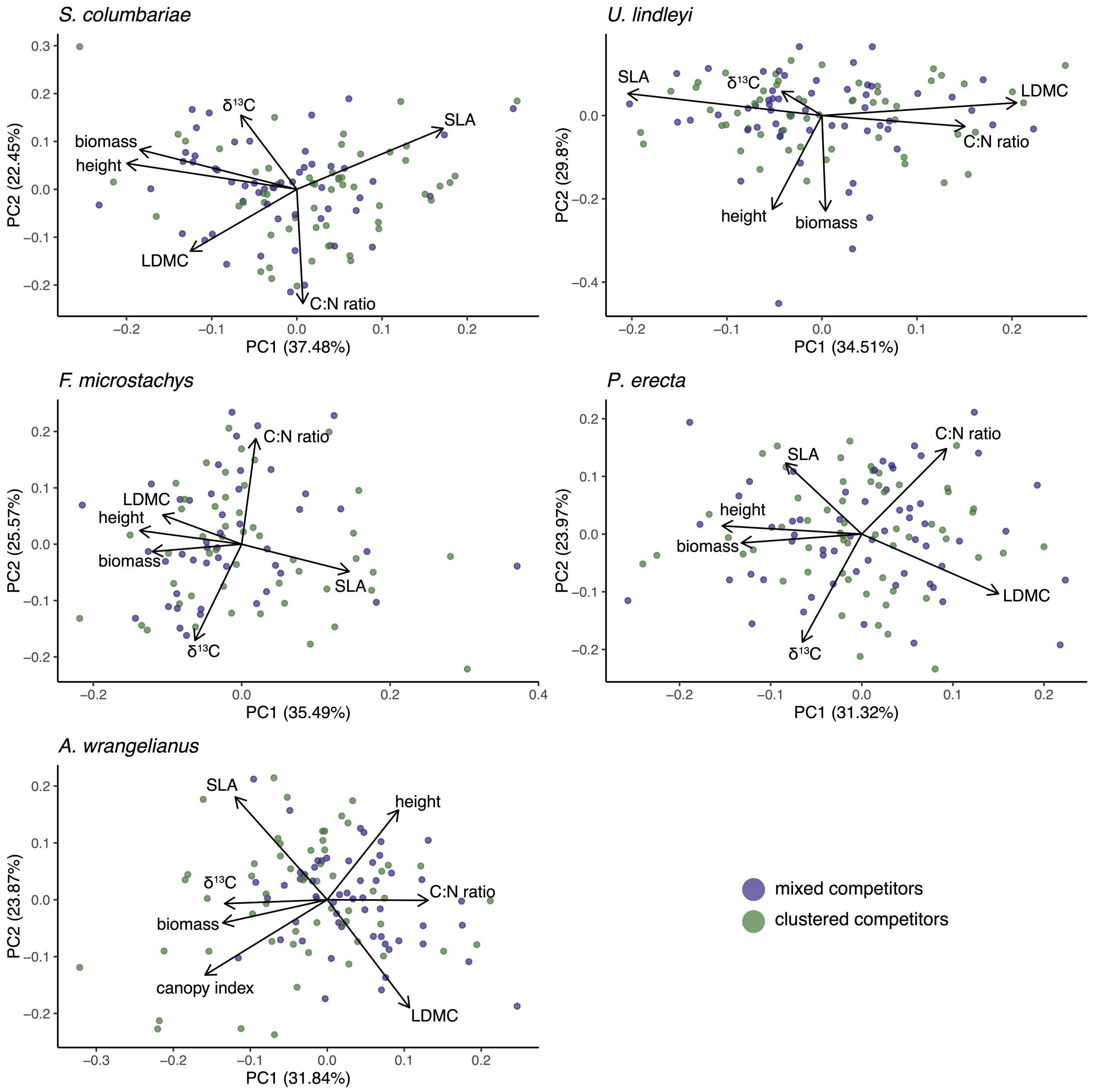

Figure S8. Principal components analysis (PCA) for each background competitor species. The first and second principal components (PC1 and PC2, respectively) are shown on the axes with the variance captured. Loadings are labeled with the performance metrics and functional traits we included in our analysis.

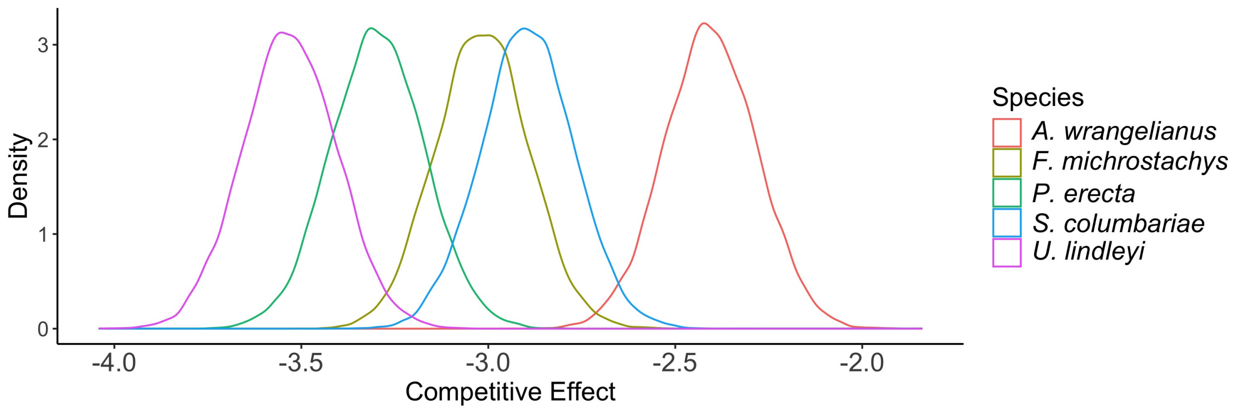

Figure S9. Posterior distributions for the competitive effect of each background competitor summed across the two spatial treatments. More negative values indicate a larger competitive effect. Colors indicate the identity of the competitor.

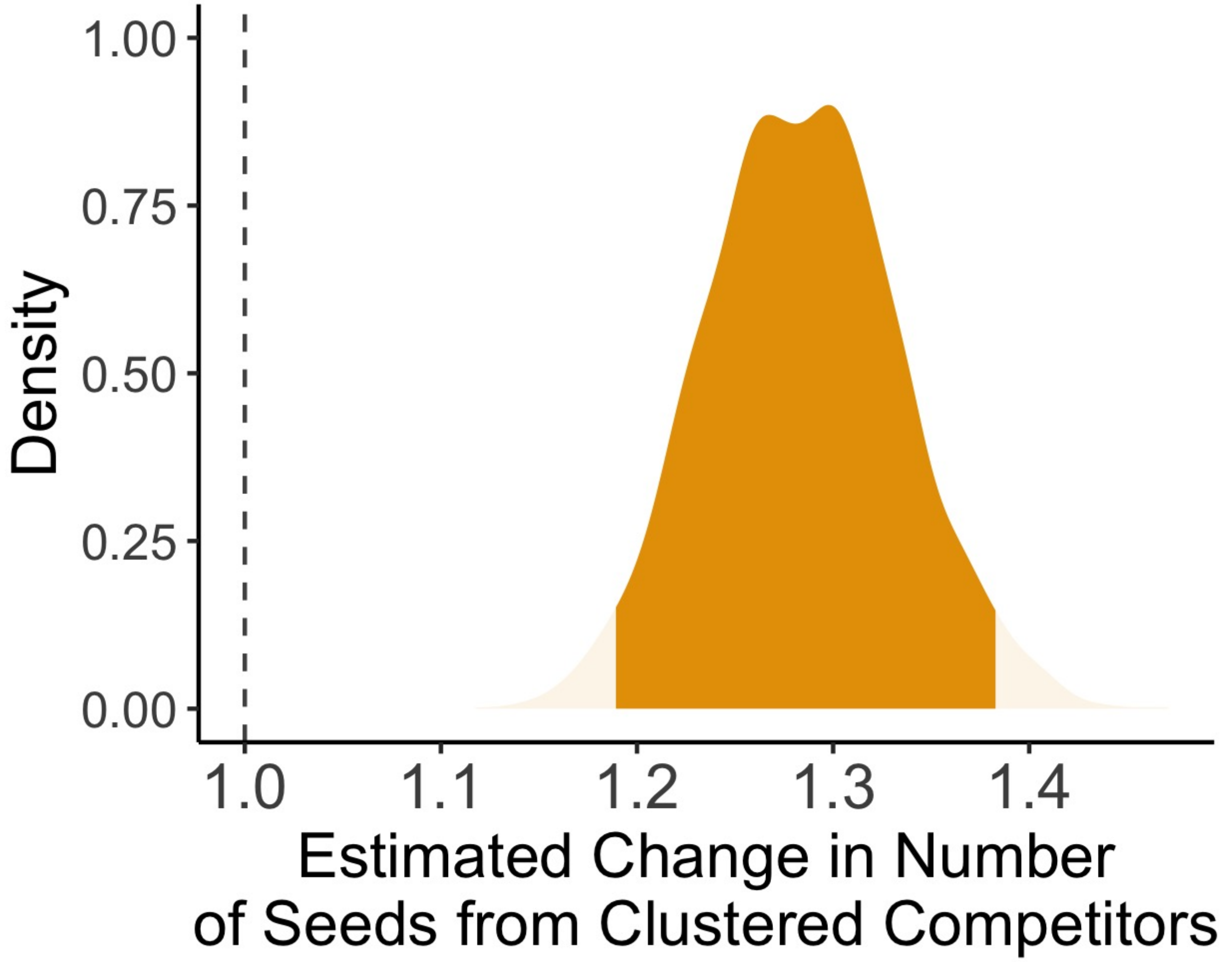

Figure S10. Posterior density for the estimated change in seed production from focal individuals in clustered, rather than mixed plots. Densities are semi-transparent outside of the $95$-th percentile of the distributions. Orange colors denote positive values while blue colors denote negative ones. We modeled seed production as a log-normally distributed random variable, fit fixed effects for focal species identity, background competitor combination and spatial arrangement.

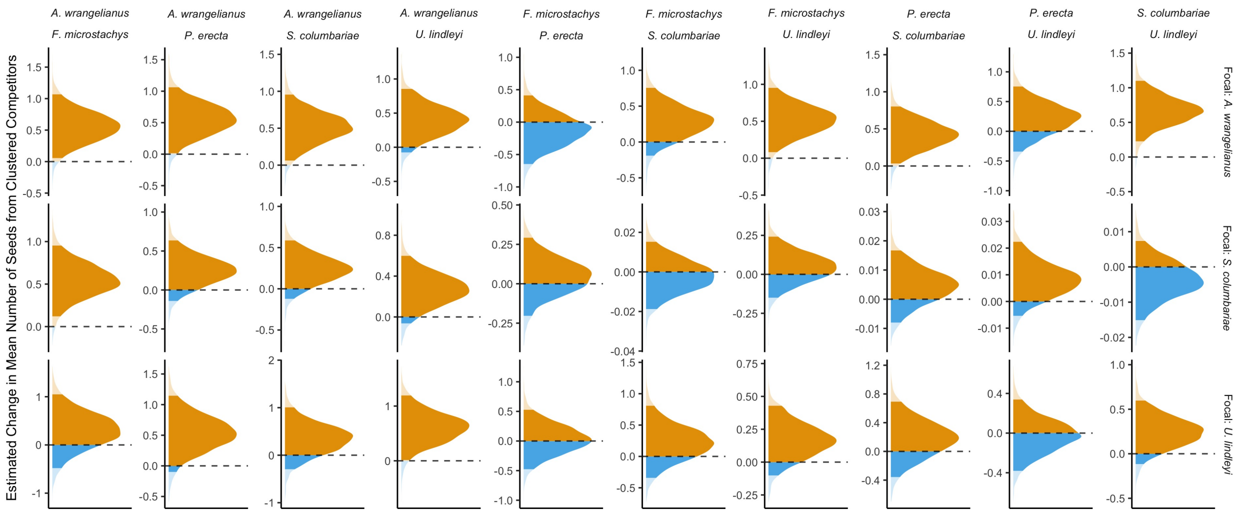

Figure S11. Posterior densities for the effect of spatial arrangement on focal seed production using a different model for each focal species (rows) and background competitor combination (columns). Otherwise, these graphs (and statistical models) are the analogous to those presented in Fig. 2 of the main text.

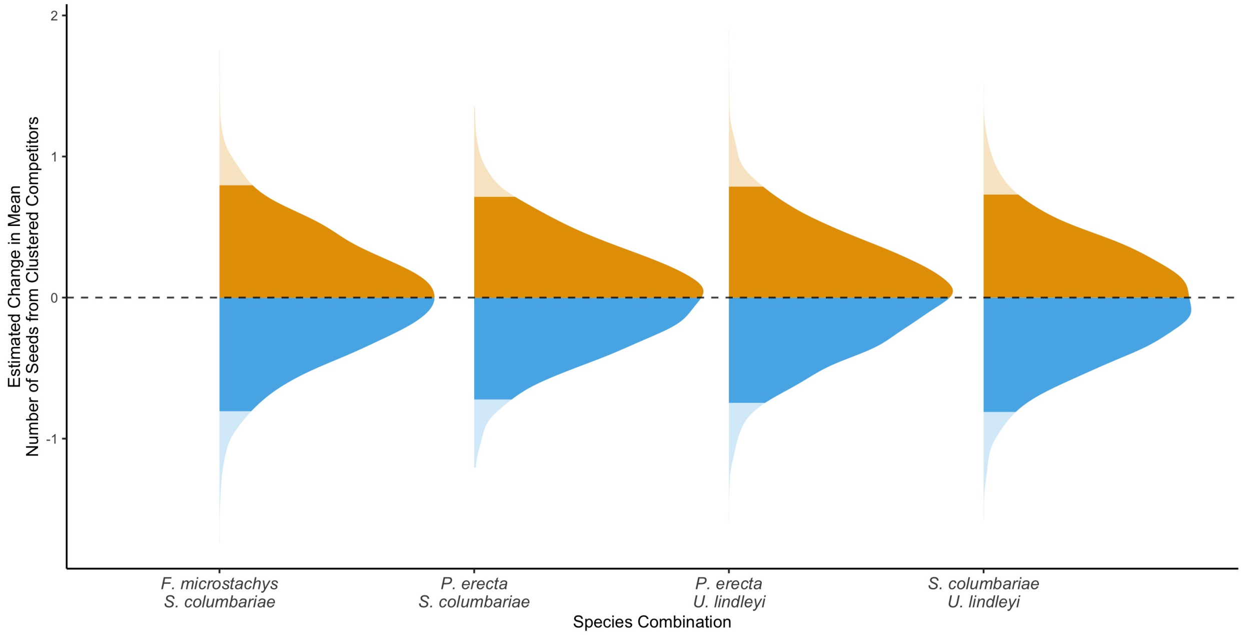

Figure S12. Analogous to Fig. 2 of the main text but only for *S. columbariae* focal individuals and the background competitor combinations with tightly concentrated posterior densities in Fig. S11. In this graph, we instead model seed production as a Poisson random variable.

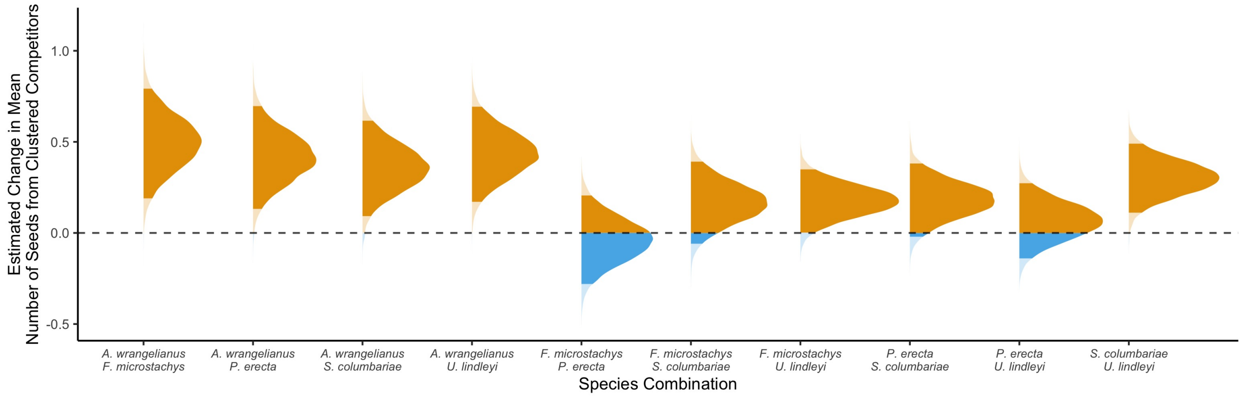

Figure S13. Analogous to Fig. 2 of the main text but modeling focal species identity as a random effect.

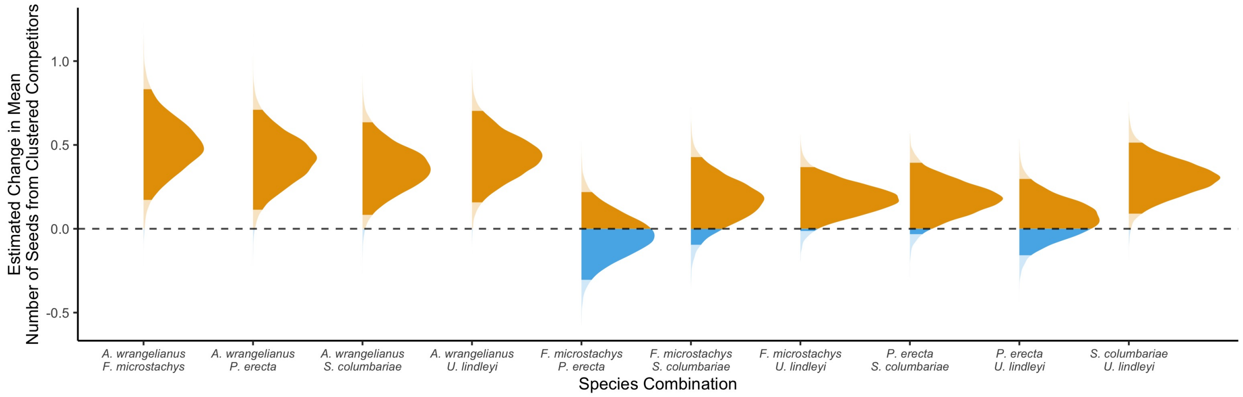

Figure S14. Analogous to Fig. 2 of the main text but including the plot identity as a random effect.

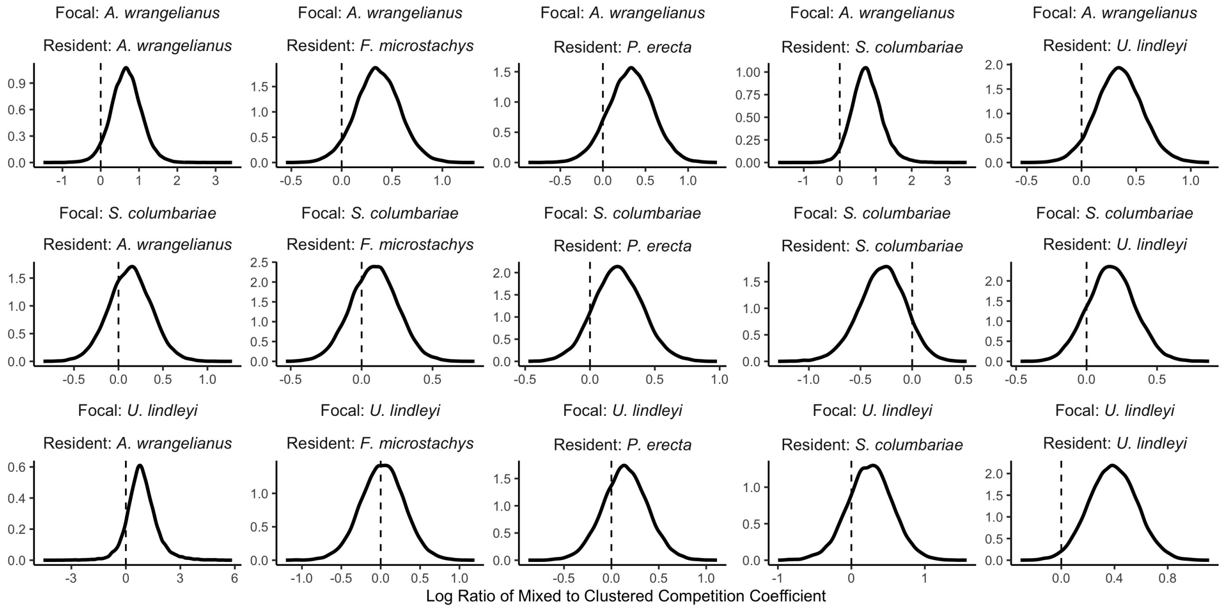

Figure S15. Posterior densities for the log ratio of competition coefficients in the Beverton-Holt model in mixed to clustered environments. Rows are different focal species and columns are different background competitors. Dashed vertical lines indicate no effect of spatial arrangement.

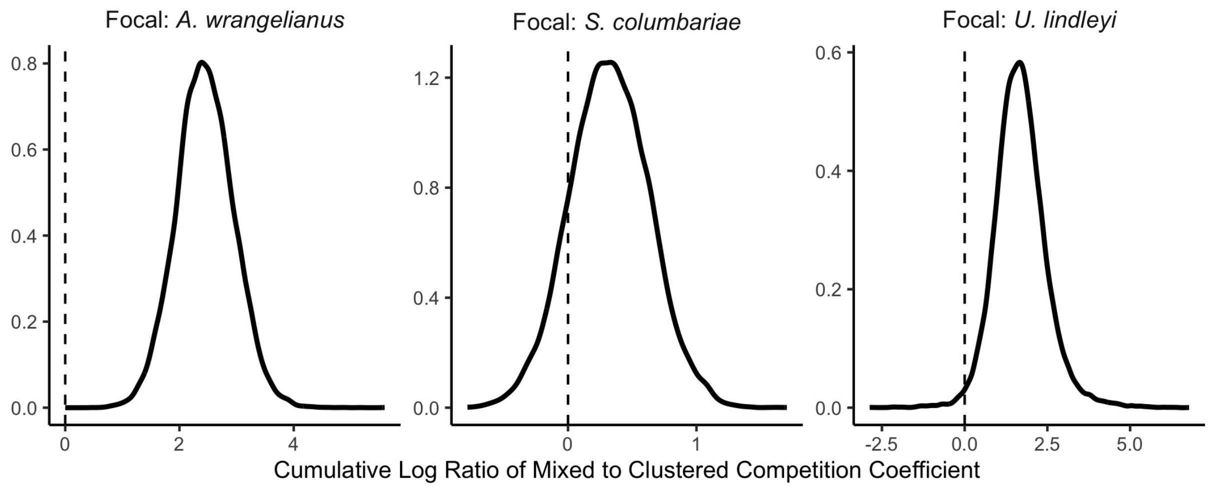

Figure S16. Posterior densities for the cumulative log ratio quantity $effect_i = \sum_j \log\left(\frac{\alpha_{ij}^{mixed}}{\alpha_{ij}^{clustered}}\right)$ which measures the effect of the spatial treatments for each focal species (panels) across background competitor combinations. Dashed vertical lines indicate no effect of spatial arrangement.

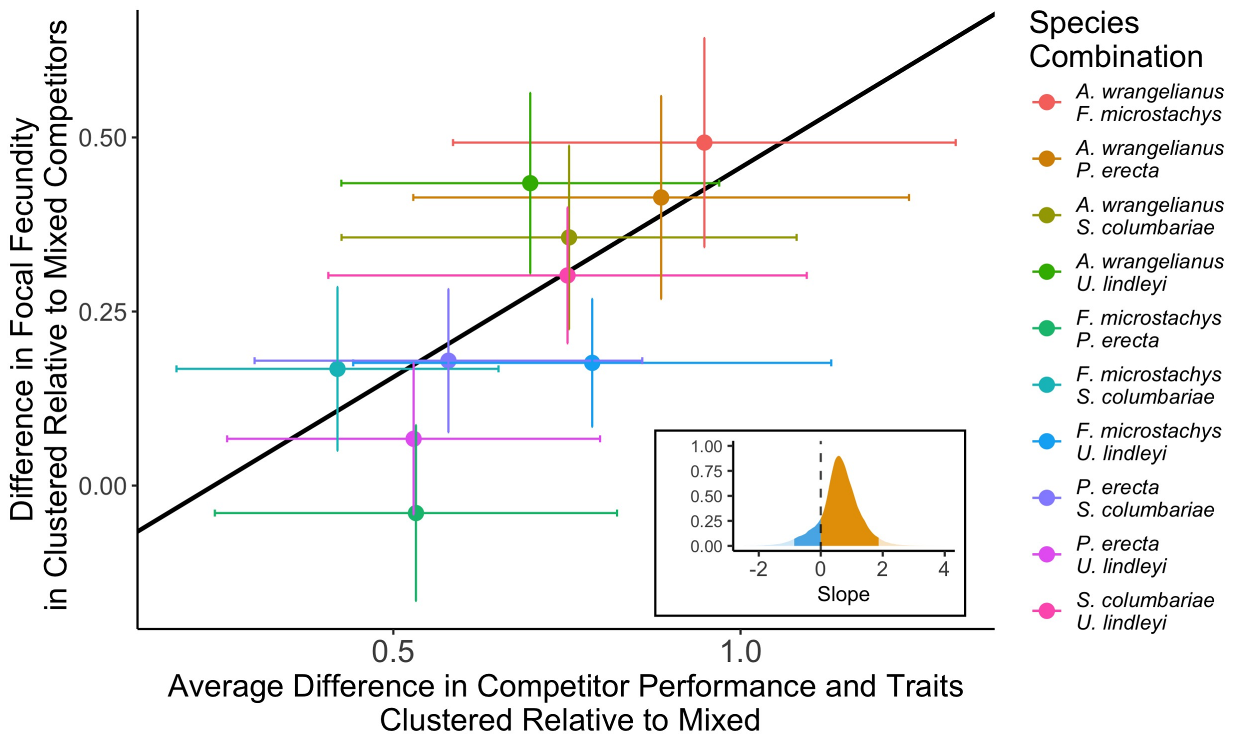

Figure S17. Analagous to Fig. 3 of the main text but using the average of the two competitor species instead of the maximum difference in the first principal component for clustered versus mixed competitors.
